# 3D micro-environment regulates NF–κβ dependent adhesion to induce monocyte differentiation

**DOI:** 10.1101/279943

**Authors:** Anindita Bhattacharya, Mahesh Agarwal, Rachita Mukherjee, Prosenjit Sen, Deepak Kumar Sinha

## Abstract

Differentiation of monocytes entails their relocation from blood to the tissue, hence accompanied by an altered physicochemical micro-environment. While the mechanism by which the biochemical make-up of the micro-environment induces differentiation is known, the fluid-like to gel-like transition in the physical micro-environment is not well understood. Monocytes maintain non-adherent state to prevent differentiation. We establish that irrespective of the chemical makeup, a 3D gel-like micro-environment induces a positive-feedback loop of adhesion-MAPK-NF-κβ activation to facilitate differentiation. In 2D fluid-like micro-environment, adhesion alone is capable of inducing differentiation via the same positive-feedback signalling. Chemical inducer treatment in fluid-like micro-environment, increases the propensity of monocyte adhesion via a brief pulse of p-MAPK. The adhesion subsequently elicit differentiation, establishing that adhesion is both necessary and sufficient to induce differentiation in 2D/3D micro-environment. Our findings challenge the notion that adhesion is a result of monocyte differentiation. Rather it’s the adhesion which triggers the differentiation of monocytes. MAPK, and NF-κβ being key molecules of multiple signaling pathways, we hypothesize that biochemically inert 3D gel-like micro-environment would also influence other cellular functions.

**Summary statement:** This article brings out a new insight into the novel mechanisms of monocyte differentiation solely driven by physical micro-environment and adhesion.

## Introduction

Cellular functions are controlled by combinations of autonomous and non-autonomous factors. Non-autonomous factors elicit cellular responses via the cellular micro-environment. The chemical makeup of a cellular micro-environment imparts its response either via the receptor-ligand interaction at the plasma membrane or through the transport of soluble molecules into the cytoplasm. The physical makeup of the micro-environment also influences the functioning of the cells^1^ ^2^ ^3^ ^4^. The functional heterogeneity in the macrophages associated with different tissues is speculated to arise from their heterogeneous micro-environments^5^ ^6^. Inside an organism, cells encounter three distinct type of physical micro-environments i) 3D fluid-like (e.g nonadherent cells in the blood), ii) 2D fluid-like (e.g endothelial, adherent cells covered by fluid) and iii) 3D gel-like (e.g adherent cells deep in the tissue). The mechanism by which the physical makeup of a micro-environment imparts cellular response is poorly understood. Monocytes provide an ideal model system to investigate such questions, for a monocyte experiences, distinct physical micro-environments during its journey to become a macrophage^7^ ^8^.

While the yolk sac-derived (embryonic origin) tissue-resident macrophages^9^ are maintained by self-renewal, the blood-derived (hematopoietic origin)^10^ ^5^ macrophages are maintained by a continuous supply of monocytes from the blood. The non-adherent monocytes migrate continuously from the blood (a complex fluid^11^) to the tissue (a viscoelastic gel^12^ ^13^ ^14^), to get differentiated into adherent macrophages. *In vitro* experiments with monocyte differentiation involves chemical stimulation on a glass/plastic substrate. Although such experiments mimic the change in chemical micro-environment faithfully, they fail to imitate the changes in the physical micro-environments. In this manuscript, we explore the influence of 3D gel-like micro-environment on monocytes (THP-1/HL-60/Peripheral blood-derived monocyte cells (PBMC)). Our results indicate that a suitable physical micro-environment is self-sufficient to trigger monocyte differentiation even in absence of chemical inducers. We also identify the signaling networks that are responsible for differentiation.

## Results

### Chemical stimulation of monocytes gives rise to ‘adherent’ and ‘non-adherent’ cell subpopulations

Chemical stimulation of THP-1 cells with established chemical inducers^15^ such as Lipopolysaccharide (LPS) or Phorbol 12-myristate 13-acetate (PMA) in an adhesion compliant Petri dish, differentiate them into macrophages. Fig. 1A depicts the Western blot and RT-PCR analysis of THP-1 cells for monocyte (CD35)^16^ ^17^ ^18^ and macrophage (CD68) markers^16^ ^18^. Non-adherent THP-1 cells (**NADH-THP-1**) when cultured in an adhesion compliant Petri dish for 5 days; a small fraction of them spontaneously adhere to generate a subpopulation of adhered cells (**ADH-THP-1**) (Fig. 1B).While the NADH-THP-1 continues to be CD35 positive, the ADH-THP-1 stops expressing CD35 and become CD68 positive (Fig. 1B). Chemical induction of NADH-THP-1 cells with LPS/PMA in an adhesion compliant Petri dish increases the fraction of ADH-THP-1 cells compared to control (Fig. 1C and Fig. S1A). We also observe that a significant fraction of chemically stimulated THP-1 cells fail to adhere irrespective of dose and duration of the inducer treatment (Fig. 1C and Fig. S1A). This confirms that irrespective of inducer treatment the ADH-THP-1 cells differentiate to macrophages whereas the NADH-THP-1 cells continue to be monocytes.

**Fig. 1.**
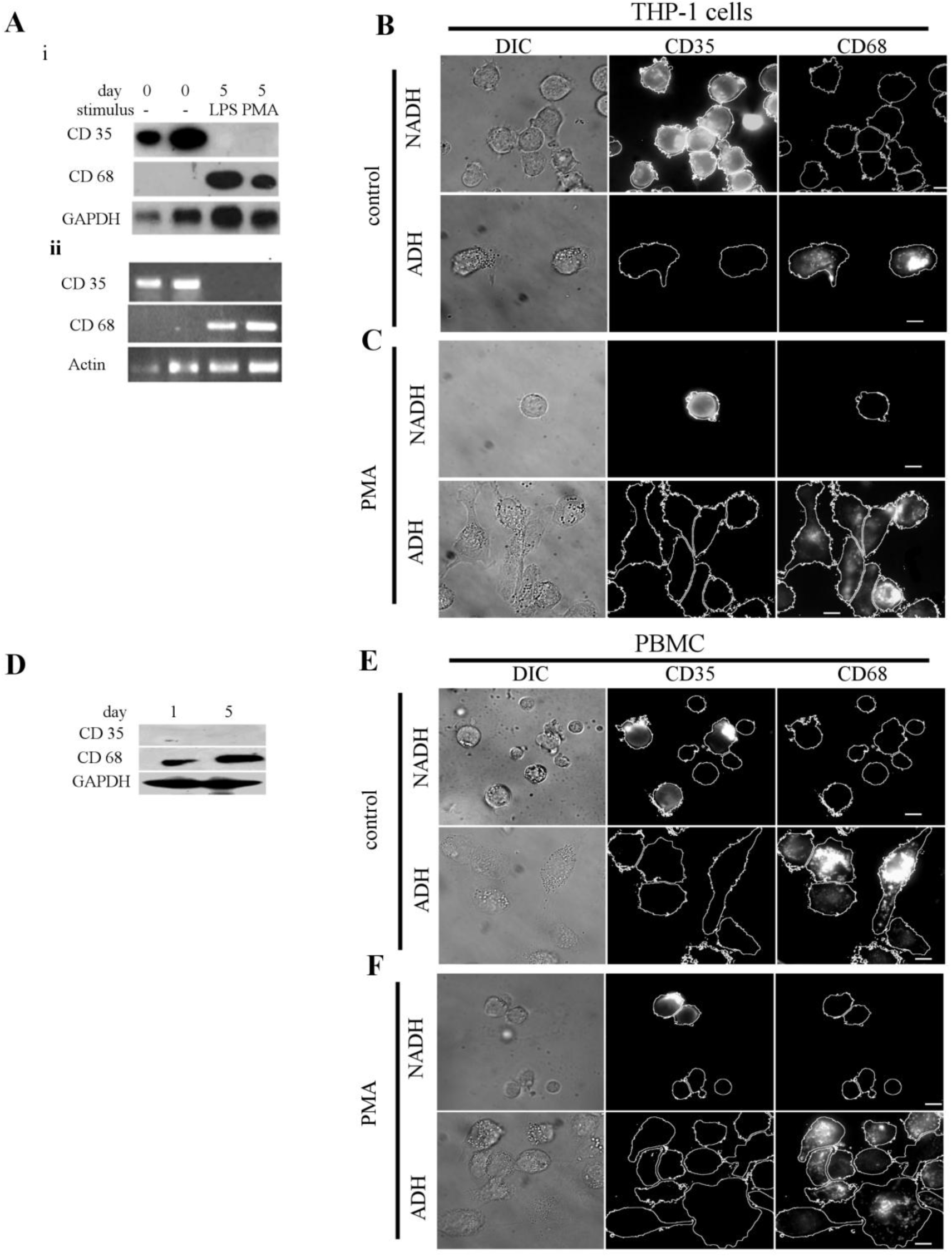
Chemical stimulation of monocyte cells gives rise to two populations of cells with distinct phenotypes. (A) (Ai) Western blot and (Aii) semi-quantitative RT-PCR of CD35 and CD68 markers in THP-1 cells, the increased GAPDH/Actin in lane 2 is because of the proliferation of monocytes and in lane 3 and 4 because of increased cellular size. (B-C) DIC (left), immunostain of CD35 (middle) and immunostain of CD68 (right) images of the control (B) and PMA treated (C) non-adherent (NADH-THP-1) (upper panel) and adherent (ADH-THP-1) (lower panel) fraction of THP-1 cells. The images in ‘B’ and ‘C’ are taken on 5^th^ day post induction/seeding. (D) Western blot of CD35 and CD68 markers in PBMC cells. (E-F) DIC (left), immunostain of CD35 (middle) and immunostain of CD68 (right) images of the control(E) and PMA treated (F) non-adherent (NADH-PBMC) (upper panel) and adherent (ADH-PBMC) (lower panel) fraction of PBMC cells. The images in ‘E’ and ‘F’ are taken on 5^th^ day post induction/seeding. (Scale bar 10 µm)

Since PBMC is already induced by cytokines/interleukins in the blood^19^ ^20^ ^21^, therefore, we get low levels of CD68 on day 1and observe a significant increase of CD68 on day 5 (Fig. 1D) post isolation. Immunostaining reveals two sub-populations of PBMC, ‘adherent’ (CD68; Fig. 1E) and ‘non-adherent’ (CD35; Fig. 1F). This establishes that PBMC and THP-1 exhibit identical behaviour with regard to their relation between adhesion and differentiation.

### Spontaneously adhered monocytes phenotypically resemble macrophages obtained by chemical induction

Fig. 2A revalidates the arrest of cell cycle at a G0/G1 stage in LPS/PMA treated THP-1 cells^22^. Next, we investigated if both the sub-populations of inducer treated NADH-THP-1 and ADH-THP-1 cells undergo cell-cycle arrest. For this, we used single-cell proliferation assay (SCPA). Fig. 2B depicts the representative images of PMA treated single THP-1 cell in 96-well-plate on day-0 and day-5. While some of the stimulated single THP-1 cells get G0/G1 arrested and remain single cell (Fig. 2Bii), the other single cells continue to proliferate and give rise to a colony of cells by day 5 (Fig. 2Biii). Incidentally, all the non-proliferating cells in different wells on day 5 are adherent (ADH-THP-1) whereas all the proliferating cells in different wells are non-adherent (NADH-THP-1). We reconfirmed the differentiation of ADH-THP-1 in SCPA by immunostaining (Fig. 2C). The NADH-THP-1 cells in SCPA after 15 days of proliferation are observed to be monocytes (CD35 positive, Fig. 2D). Thus the chemically induced NADH-THP-1 cells do not undergo cell cycle arrest. We observed (Fig. 2E-F) that even after prolonged incubation at a considerably higher concentration of PMA, a significant fraction of THP-1 cells continues to be monocyte (NADH-THP-1). Next, we investigated the phagocytotic ability of ADH-THP-1 and NADH-THP-1 cells ^23^. Fig. 2G, H confirm that irrespective of chemical inducer treatment only a fraction of ADH-THP-1 are phagocytotic (Fig. S2B,C) and none of the NADH-THP-1 are capable of phagocytosis. As observed with ADH-THP-1, only a fraction of PBMC are phagocytotic (Fig. 2G).

**Fig. 2.**
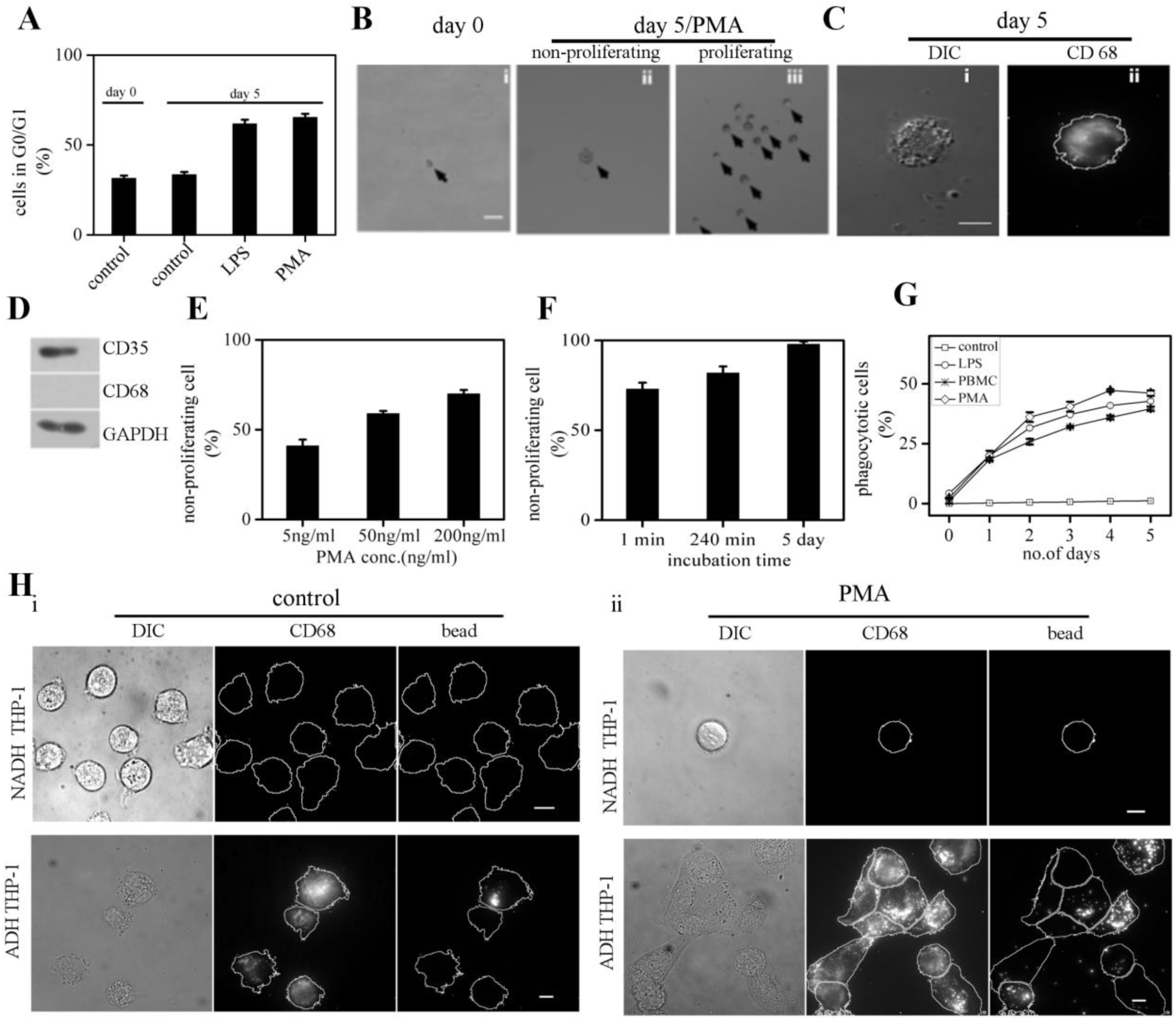
An adherent subpopulation of cells functionally resemble macrophages. (A) Plot depicting the percentage of G0/G1 arrested cells in control and LPS/PMA treated THP-1 cells. (B), Phase contrast images of PMA treated single THP-1 cell on day 0 (Bi), and day 5 (Bii,Biii). ‘Bii’ and ‘Biii’ depicts non-proliferating and proliferating PMA treated cells respectively. (C) DIC (Ci) and immunostain of CD68 (Cii) images of non-proliferating single THP-1 cell (ADH-THP-1) on 5^th^ day post PMA induction. (D) Western blot of CD35 and CD68 in PMA induced proliferating THP-1 cells. (E-F) Plot depicting the percentage of non-proliferating THP-1 cells for different doses (E) and durations (F) of PMA induction. (G) Plot depicting the percentage of phagocytotic THP-1 and PBMC cells. (H) DIC (Left) immunostain of CD68 (middle) and phagocytised fluorescent beads (right) images of the control (Hi) and PMA treated THP-cells (Hii). The non adherent (NADH-THP-1) and adherent (ADH-THP-1) fraction of cells are depicted in upper and lower panel respectively. (Scale bar 50 µm for b and 10 µm for c and h)

Thus our observations confirm that adhesion of monocytes and their differentiation into macrophage are inter-dependent. Therefore, next we dissected the effect of adhesion and chemical induction on monocytes with regard to their differentiation.

Adhesion is necessary for the survival of many adherent cells types^24^ ^25^ ^26^. However, whether adhesion is essential for viability of ADH-THP-1 is not known. We stimulate the THP-1 cells in conditions which are incompatible for cellular adhesion (Fig. 3A-C) MTT absorbance confirms the viability of stimulated THP-1 cells without adhesion for 5 days (Fig. 3D, Fig. S3Aa,B). We also observe the formation of cellular clumps of stimulated THP-1 (ADH-THP-1) cells (Fig. 3B) due to overexpression of E-cadherins in adhesion incompatible conditions (Fig. 3E)

**Fig. 3.**
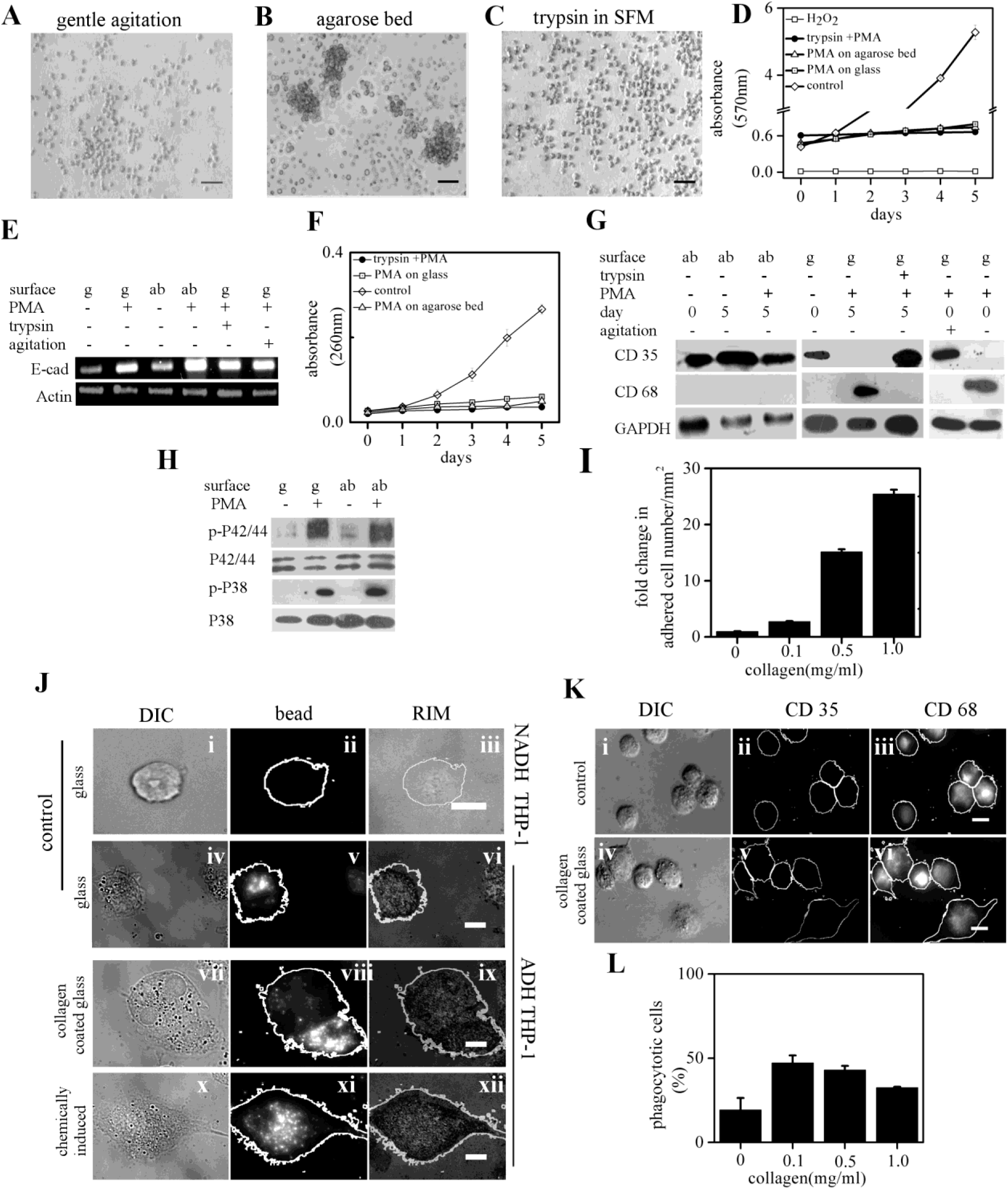
Adhesion is both necessary and sufficient for monocyte to macrophage differentiation in 2D culture. (A-C) Phase contrast images of PMA treated THP-1 cells under periodic agitation on glass surface (A), on agarose bed (B) and on coverglass in serum-free media with 0.25% trypsin (C). (D) MTT absorbance at 570 nm obtained from lysate of THP-1 cells cultured under different adhesion conditions on different days post PMA induction. (E) Semi-quantitative RT-PCR of E-cadherin of THP-1 cells cultured in different adhesion conditions. (F) Genomic DNA concentration (absorbance at 260 nm) from lysate of THP-1 cells cultured under different adhesion conditions on different days post PMA induction. (G) Western blot analysis for CD35 and CD68 on THP-1 cells under different adhesion conditions. Experiments with gentle agitation was analysed 8 hr post seeding/induction on day 0. (H) Western blot analysis of p-P42/44 MAPK, p-P38 MAPK of THP-1 cells under different conditions 15 min post PMA induction. (I) Plot depicting the density of spontaneously adhered THP-1 cells on collagen-coated (0-1 mg/ml) glass substrate. (J) DIC (left), phagocytised fluorescence beads (middle) and RIM (right) images of non-adherent (Ji-iii) and adherent (Jiv-xii) THP-1 cells on day 5 post seeding/induction. (K) DIC (left), immunostain of CD35 (middle), and immunostain of CD 68 (right) images of THP-1 cells on day 5 post seeding. (L) The percentage of phagocytotic THP-1 cells seeded on collagen-coated coverglass (0-1 mg/ml). (ab: agarose bed, g: glass. Scale bar 50µm for a, b, c and 10 µm for j,k),

### Adhesion is a necessary biophysical condition for differentiation of THP-1 cells in 2-D culture

We observe a gradual increase in the mitochondrial activity (Fig. 3d) and genomic DNA concentration (Fig. 3F) in the THP-1 cells caused by their proliferation. A significant reduction of cell proliferation caused due to PMA/LPS treatment in adhesion incompatible conditions (Fig. 3D.F) suggests its cell cycle arrest. Though the cell cycle of stimulated THP-1 (other than NADH-THP-1) cells is arrested, these cells fail to complete differentiation (Fig. 3G) without adhesion. Fig. 3H confirms that early signaling events (within 15 minutes of stimulation) such as phosphorylation of MAPK ^27^ are independent of adhesion. Thus, these results suggest that adhesion is a necessary biophysical condition for completing the entire process of monocyte differentiation after chemical stimulation.

### Adhesion is sufficient for the differentiation of THP-1 cells in 2D culture

Next, we investigated the sufficiency of adhesion in setting off the process of differentiation. We observe that THP-1 cells in adhesion promoting conditions (collagen coating; 0-1 mg/ml) in the absence of any chemical inducer give rise to a significantly higher fraction of ADH-THP-1 (Fig. 3I). In reflection interference microscopy (RIM), the regions of the plasma membrane which are closer to the glass appear dark because of destructive interference ^28^ ^29^. Using this we find comparable adhesions between inducer free and inducer activated ADH-THP-1 (Fig. 3J). The RIM patches appear to be equally dark under all the conditions, suggesting no significant difference in the glass and plasma membrane proximity between inducer free (Fig. 3Jiv-Jix) and chemically induced ADH-THP-1 (Fig. 3Jx-Jxii) cells. Interestingly, irrespective of collagen coating, ADH-THP-1 cells express CD68 (Fig. 3K) and exhibit phagocytotic ability (Fig. 3Jii,jv,jviii, jxi, L) like that of chemically induced ADH-THP-1 cells. However, the fraction of phagocytotic cells differs from that of PMA induced ADH-THP-1 cells (Fig. 3L). A possible explanation for this difference is discussed in the supplementary appendix-1. Therefore, we conclude that adhesion is sufficient to promote differentiation in THP-1 cells irrespective of presence/absence of any chemical stimulus.

### Chemical induction alters the membrane fluctuation to facilitate adhesion in THP-1 and RAW cells

As expected, the RIM images of NADH-THP-1 cells show significantly lesser dark patch (Fig. 4Aii) compared to PMA activated THP-1 cells (Fig. 4Bii). While the left cell in Fig. 4Bii does not develop stable adhesion, the cell on the right side has developed a stable adhesion as indicated by the central dark patch. Thus inducer treatment cause gradual increase in fraction of dark pixels per cell in RIM image of THP-1 cells (Fig. 4C). Careful inspection of Fig. 4Aii (inset), reveals smaller dark puncta. A time lapse study of these dark puncta reveals the transient nature of membrane and glass contact in NADH-THP-1 cells (Upper kymograph; Fig. 4D inset, Fig. S4A and video 1). PMA treatment increases the dwell time of dark puncta (Fig. 4D, video 2 and 3) probably by altering the cortex/plasma-membrane of NADH-THP-1 cells. We hypothesize that the increased duration of dark puncta must facilitate the formation of stable focal adhesion. This was confirmed with RAW cells because of its better transfectability than THP-1. Fig. 4E-G depicts the progression of adhesion post seeding in RAW cells in absence/presence of PMA. The dwell time of dark puncta (Fig. 4H, Fig. S4B and Video 4-6) in RAW cells increases upon PMA treatment. We observe the higher turnover rate of focal adhesion complex in control RAW cells. The PMA treatment reduces the turnover of focal adhesion complex as measured by its dwell time (Fig. 4I-J, Fig. S4C, and Video 7-8). Thus PMA treatment has an immediate effect such that the adhesion of the cells get stronger/stable (Fig. S4A-C).

**Fig. 4.**
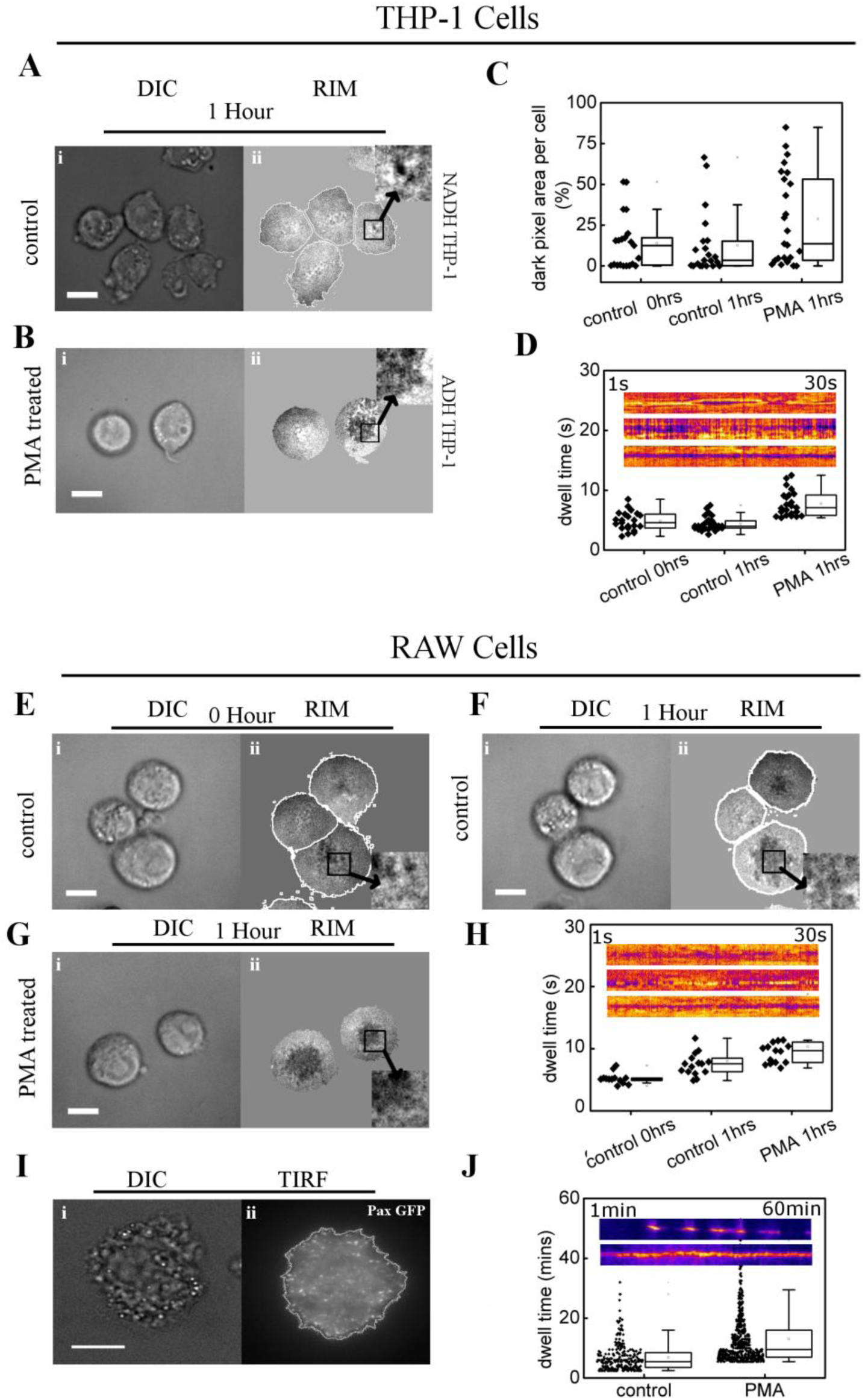
Chemical inducer alters membrane fluctuation to facilitate adhesion. (A-B) DIC (Ai, Bi) and RIM (Aii, Bii) images of control (A) and PMA treated (B) THP-1 cells 1 hr post seeding/induction (insets in aii and bii depicts the punctuated nature of dark pixels). (C) Quantification of percentage of dark pixels in RIM images of THP-1 cells under different conditions. (D) The dwell time of dark puncta (inset of aii, bii) in control and PMA treated THP-1 cells. Inset in ‘d’ depicts Kymograph of dark puncta in control (0 hrs post seeding; upper), control (1hr post seeding; middle) and PMA treated (1 hr post induction; lower) cells. (E-G) DIC (Ei, Fi, Gi) and RIM (Eii, Fii, Gii) images of control (0 hr post seeding (E) and 1 hr post seeding (F), and PMA treated (1 hr post induction /seeding (G) RAW cells. (insets in Eii, Fii and Gii depicts the punctuated nature of dark pixels). (H) The dwell time of dark punta (inset of Eii Fii and Gii) in control(0 hr post seeding), control(1 hr post seeding) and PMA induced(1 hr post induction /seeding) cells. Inset in ‘H’ depicts Kymograph of dark puncta in control (0 hrs post seeding; upper), control (1hr post seeding; middle) and PMA treated (1 hr post induction; lower) cells. (I) DIC (i) and TIRF (ii) images of PMA treated RAW cells expressing Paxillin-GFP. (J) The dwell time of focal adhesion in control (1 hr post seeding) and PMA treated (1 hr post induction/seeding) RAW cells. Inset in ‘J’ depicts Kymograph of focal adhesion in control (1 hrs post seeding; upper) and PMA treated (1 hr post induction; lower) cells. (Scale bar 10 µm)

### Adhesion of THP-1 cells activate NF-κβ which mediates monocyte differentiation

Independent literature survey suggests NF-κβ activation is associated with both cellular adhesion ^30^ ^31^ and monocyte differentiation^32^ ^33^. Therefore, we hypothesize that adhesion mediated activation of NF-κβ is necessary for differentiation of monocytes as depicted in Fig. 5A. While the MAPK activation in response to chemical induction lasts only for short duration (0-30 min post-induction; Fig. 5Bi, and Fig. 3H) in absence of adhesion, the adhesion maintains a higher level of activated MAPK at later time points (Fig. 5Bi-Bii). As a result activated NF-κβ levels also depend on the adhesion (Fig. 5C). The short pulse of MAPK activation suggests the existence of negative feedback loop (as proposed in Fig. 5A). Alike chemically stimulated (LPS and PMA) populations of ADH-THP-1 cells, the spontaneously adhered ADH-THP-1 in inducer free conditions exhibit NF-κβ activation (Fig. 5Ci). This suggests that activation of NF-κβ is independent of chemical inducer treatment. This is further confirmed in Fig. 5Cii, notwithstanding inducer treatment, NF-κβ is not activated in NADH-THP-1 cells. Thus Fig. 5C confirms that adhesion activates NF-κβ in monocytes. Fig. 5D-F confirm that p-MAPK acts upstream of NF-κβ activation (Fig. 5A). The P42/44 MAPK and P38 MAPK molecules act parallel since inhibition of either of them does not lead to inhibition of the other (data not shown). Yet, simultaneous activation of P42/44 MAPK and P38 MAPK is necessary for NF-κβ activation. Fig. 5G and Fig. S5 confirm the necessity of NF-κβ and MAPK activation for monocyte differentiation. Next, we investigated the relevance of short pulse of phosphorylated MAPK (Fig. 5B) in response to chemical induction.

**Fig. 5.**
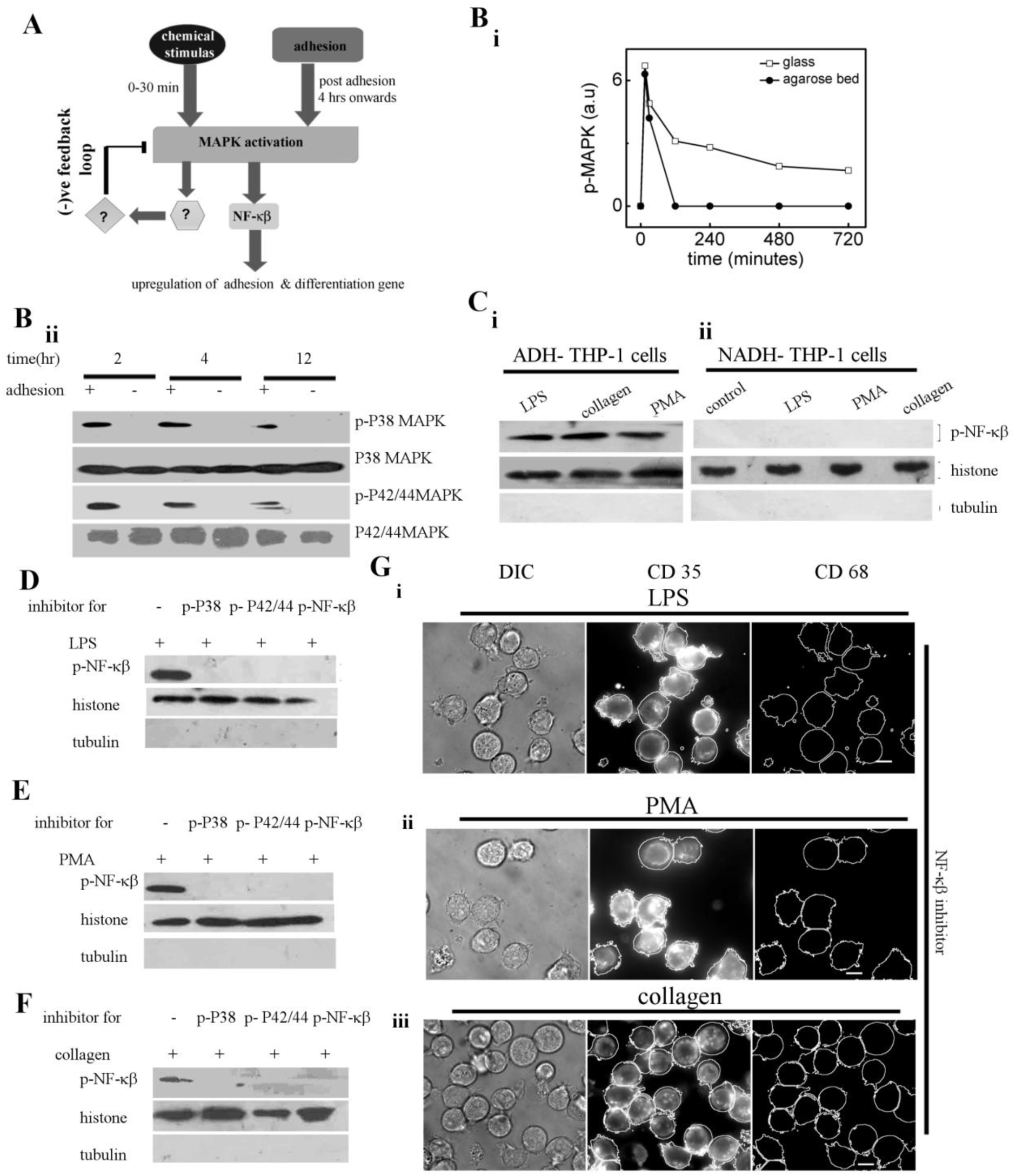
Adhesion dependent activation of NF-κβ is responsible for differentiation. (A) Schematic of signaling cascade depicting the key molecules in monocyte to macrophage differentiation. (B) Graphical representation of average of p-P38 MAPK and p-P42/44 MAPK levels at different time post induction with/without adhesion (Bi). Western blot of p-P38 MAPK and p-P42/44 MAPK of PMA stimulated THP-1 cells at different time post induction with/without adhesion (Bii). (C) Western blot of p-NF-κβ in the nucleur fraction of adherent (ADH, Ci) and non-adherent (NADH, Cii) populations of THP-1 cells in different conditions. (D-F) Western blot of p-NF-κβ in the nuclear fraction of THP-1 cells stimulated with LPS (D), PMA (E) and collagen-coated surface (F) under different conditions of inhibitor treatment (U0126: p42/44MAPK, SB203580: P38 MAPK and NF-κβI: NF-κβ). (G) DIC (left column), immunostain of CD35 (middle column) and CD 68 (right column) of THP-1 cells treated with NF-κβI under stimulation/seeding with LPS (Gi), PMA (Gii) and collagen-coated surface (Giii). (Scale bar 10 µm)

### Positive feedback loop regulates the adhesion in THP-1 cells

We hypothesize that the short pulse of activated MAPK (Fig. 3H and Fig. 5B) must be responsible for changing the adhesion state of the monocytes. THP-1 cells make stable adhesion 6-8 hrs post induction (see method, data not shown). Fig. 6A and Fig. S6 confirms that many genes involved in focal adhesions are transcriptionally up-regulated upon chemical induction prior to formation of stable focal adhesion. Interestingly, the inducer-free collagen mediated adhesion also leads to transcriptional up-regulation of adhesion genes (Fig. 6A).

**Fig. 6.**
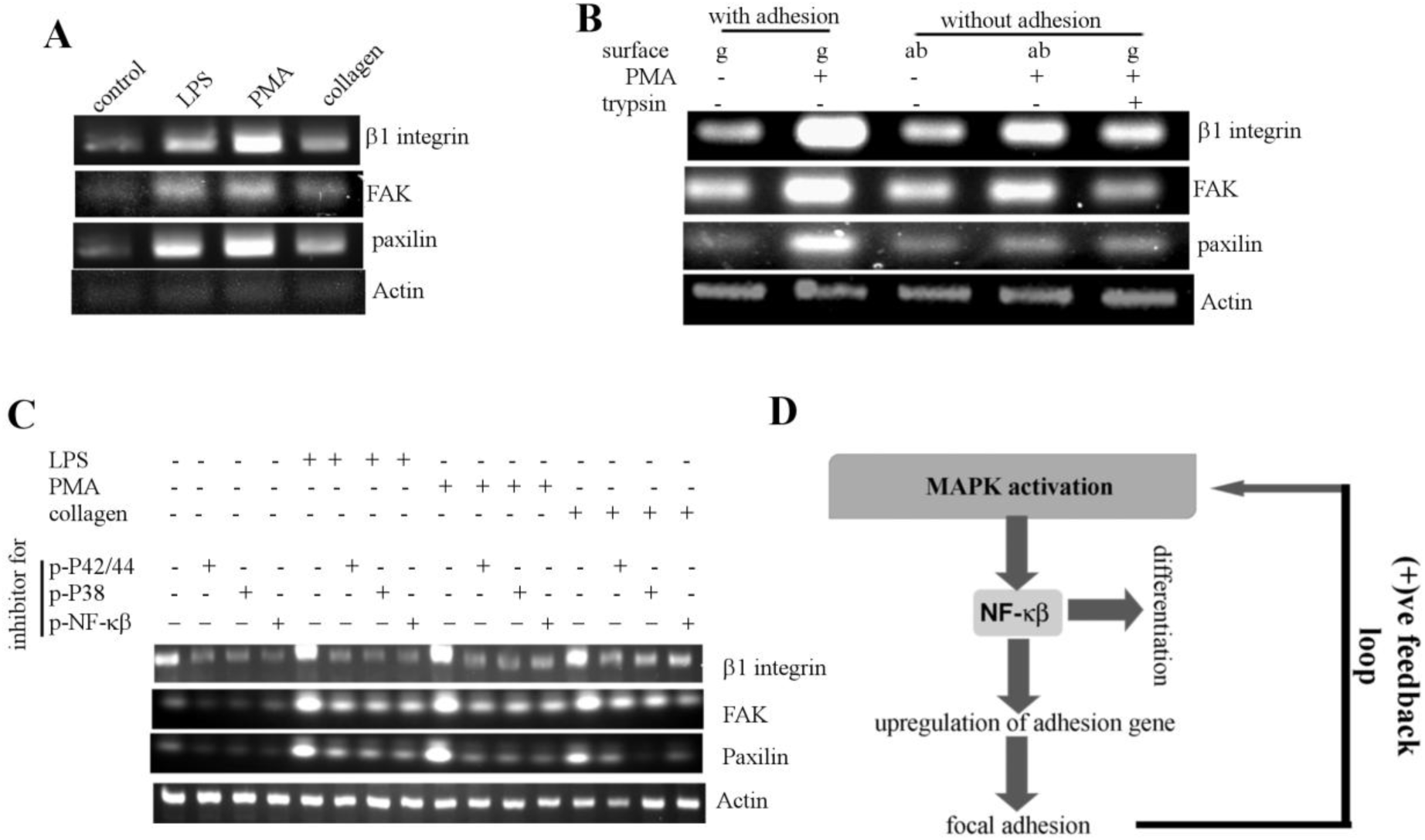
Adhesion mediated positive feedback loop regulates the p-NF-κβ levels in ADH-THP-1 cells. (A) Semi-quantitative RT-PCR of adhesion genes (β1 integrin, Paxilin and FAK) in THP-1 cells under different conditions (6 hrs post seeding/induction). (B), Semi-quantitative RT-PCR of adhesion genes in THP-1 cells under different conditions of adhesion/chemical-stimulation. (C) Semi-quantitative RT-PCR of adhesion genes in THP-1 cells under different conditions of chemical-stimulation in presence/absence of p-P42/44MAPK (U0126), p-P38MAPK (SB203580) and NF-κβ (NF-κβI) inhibitors. (D) Schematic of signalling cascade depicting the positive feedback loop of adhesion which leads to monocyte to macrophage differentiation

Therefore, we hypothesize that an adhesion driven positive feedback loop exists for changing the adhesion status of the monocytes. Fig. 6B (compare lane 1with 4 and 1 with 5) confirms transcriptional up-regulation of the adhesion genes in response to inducer treatment without adhesion. Allowing adhesion to chemically induced THP-1 leads to much higher levels of transcriptional up-regulation of the adhesion genes (compare lane 2 with 4 and 2 with 5) confirming the existence of positive feedback loop. Fig. 6C establishes that inhibition of either of MAPK or NF-κβ prevents the transcriptional up-regulation of adhesion-related genes, suggesting MAPK and NF-κβ dependent up-regulation of these genes. Thus, we establish that a negative-feedback loop and an adhesion-dependent positive feedback loop involving MAPK activation operate simultaneously to achieve differentiation, as indicated in Fig. 6D and Fig. 5A. Having explored the link between adhesion and differentiation, next we investigated the effect of 3D gel like micro-environment on monocytes.

### 3D gel-like micro-environment of THP-1 cells induces spontaneous differentiation

The THP-1 cells cultured on glass bottom Petri dish were provided 3D gel/liquid like micro-environment by covering it with 0.1% agarose either in liquid or gel state (Fig. 7A,B). Identical chemical but distinct physical micro-environment, elicit different responses in monocytes. While in 3D liquid like micro-environment, the THP-1 cells do not differentiate spontaneously in 5 days (Fig. 7A,C), in a 3D gel-like micro-environment the THP-1 cells differentiates to macrophages (Fig. 7B,C). We also verified that the phagocytotic ability (Fig. S7A) of these macrophages is comparable to that of ADH-THP-1 cells in 2D.

**Fig. 7.**
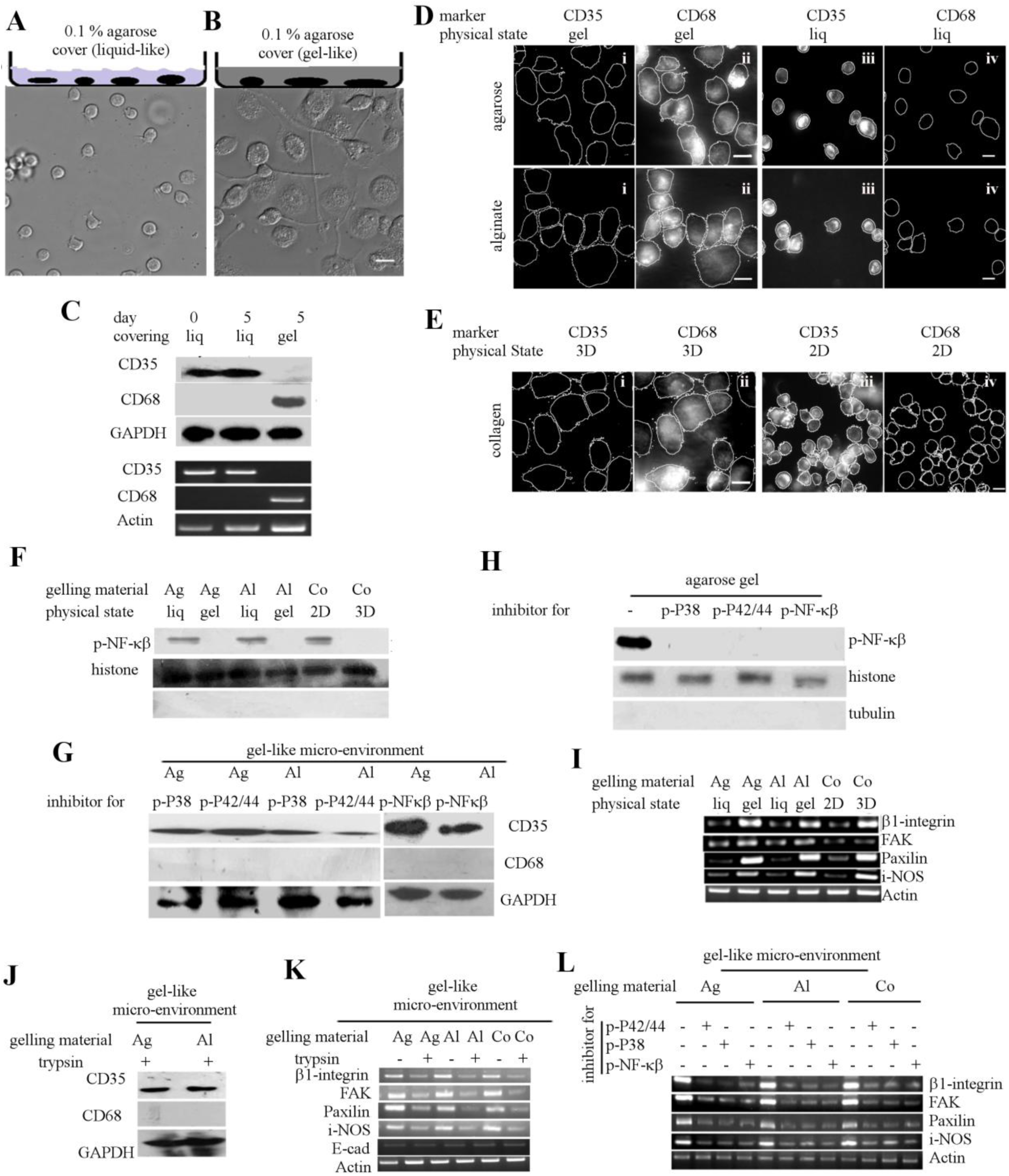
3D gel-like micro-environment induces adhesion to facilitate differentiation. (A-B) The schematic of the experiments are depicted on top of corresponding DIC images of THP-1 cells in RPMI media on day-5 covered with 0.1% agarose liquid (A) or gel (B). (C) Western blot (upper) and semi-quantitative RT-PCR (lower) analysis for CD35 and CD68 in THP-1 cells, cultured with cover of identical chemical composition (0.1% agarose) in different physical state (liq: liquid, gel: agarose gel). (D) Immunofluorescence images of THP-1 cells for CD68 and CD35 cultured in different micro-environment (gel like or liquid like cover of agarose and alginate).(E) Immunofluorescence images for CD68 and CD35 of THP-1 cells cultured in 2D or 3D collagen matrix. (F) Western blot of p-NF-κβ in the nuclear fraction of THP-1 cells isolated from different liquid-like or gel-like micro-environment of alginate (Al) and agarose (Ag). Western blot of p-NF-κβ in the nuclear fraction of THP-1 cells isolated from 2D and 3D collagen (Co) matrix is also depicted in ‘f’.(G) Western blot of CD35 and CD68 from THP-1 cells isolated from gel of agarose (Ag) and alginate (Al) treated with SB203580 (p-P38 MAPK), U0126 (p-P42/44 MAPK) and NF-κβI (NF-κβ nuclear translocation) inhibitors. (H) Western blot of p-NF-κβ in the nuclear fraction of THP-1 cells isolated from agarose gel treated with SB203580 (p-P38 MAPK), U0126 (p-P42/44 MAPK) and NF-κβI (NF-κβ nuclear translocation) inhibitors. (I) Semi-quantitative RT-PCR of i-NOS and adhesion genes β1-Integrin, Paxillin and FAK isolated from THP-1 cells cultured in 1% alginate (Al), 0.1% agarose (Ag) in liquid (liq) or gel state. Semi-quantitative RT-PCR of i-NOS and adhesion genes β1-Integrin, Paxillin and FAK isolated from THP-1 cells cultured in 2D or 3D collagen matrix is also depicted in ‘i’. (J) Western blot analysis of CD35 and CD68 in THP-1 cells treated with trypsin in serum free conditions and cultured in agarose(Ag)/alginate(Al) gel. (K) Semi-quantitative RT-PCR of i-NOS, E-cadherin (E-cad) and adhesion genes β1-Integrin, Paxillin and FAK isolated from THP-1 cells cultured in 1% alginate gel (Al), 0.1% agarose gel (Ag) and 3D collagen (Co) matrix in presence/absence of trypsin. (L) Semi-quantitative RT-PCR of i-NOS and adhesion genes β1-Integrin, Paxillin and FAK isolated from THP-1 cells treated with SB203580 (p-P38 MAPK), U0126 (p-P42/44 MAPK) and NF-κβI (NF-κβ nuclear translocation) inhibitors, and cultured in 1% alginate (Al) gel, 0.1% agarose (Ag) gel or 3D collagen matrix. (Scale bar 10 µm)

In the above experiments, the basal surface of the THP-1 cells is in contact with the glass. Next, we dissected the role of glass during differentiation of THP-1 cells covered with agarose gel. For this, the THP-1 cells were covered with agarose gel/liquid on substrate precoated with agarose gel (Fig. S7B). Interestingly, even in absence of adhesion compatible glass, the THP-1 cells undergo differentiation in a 3D gel-like micro-environment of agarose, alginate, matrigel, and collagen (Fig. 7D, E and Fig. S7D-E, G-I). The 3D gel-like micro-environment causes cell cycle arrest of THP-1 cells however fluid-like chemically identical micro-environment does not cause cell cycle arrest (Fig. S7F). While the culture of THP-1 cells on collagen-coated surface leads to a basal level of differentiation (Fig. 3I), embedding it into a 3D collagen matrix leads to a much higher level of differentiation (Fig. 7E). HL-60 cell line also exhibits similar behavior (Fig. S7J).

### 3D gel-like micro-environment induces adhesion mediated activation of NF-κβ

Fig. 7F depicts the activation of NF-κβ in THP-1 cells in a 3D gel-like micro-environment of different composition. However, fluid-like micro-environment of identical chemical composition fails to activate NF-κβ in THP-1 cells (Fig. 7F). Therefore, we hypothesize that the spontaneous differentiation of THP-1 cells in a 3D gel-like micro-environment is mediated via the same pathways as shown in Fig. 5A and Fig. 6D. As a result, inhibition of either MAPK or NF-κβ prevents its differentiation (Fig. 7G). As expected, inhibition of either of the P38 MAPK or P42/44 MAPK prevents the activation of NF-κβ in a 3D gel-like micro-environment of agarose (Fig. 7H), alginate (Fig. S7K), and collagen (Fig. S7L).

### 3D gel-like micro-environment activates positive feedback loop of adhesion to mediate NF-κβ dependent differentiation

Fig. 7I, depicts the transcriptional up-regulation of various adhesion associated gene in THP-1 cells embedded in a 3D gel of agarose, alginate, and collagen. However, fluid like micro-environment of identical chemical composition fails to activate transcription of adhesion associated genes in THP-1 cells. Therefore, we hypothesize that the 3D gel-like micro-environment activates positive feedback loop of adhesion (Fig. 6D). Above hypothesis is validated by denial of adhesion to THP-1 cells in a 3D gel-like micro-environment using trypsin. This not only abrogates the differentiation (Fig. 7J and Fig. S7M), it also prevents the transcriptional up-regulation of adhesion-related genes (Fig. 7K). Further, inhibition of either of MAPK or NF-κβ in each gelling media prevents the transcriptional up-regulation of adhesion-associated genes (Fig. 7L). These results suggest that the positive feedback loop of adhesion in Fig. 6d is activated by 3D gel-like micro-environment.

### 3D gel-like micro-environment elicit adhesion mediated activation of NF-κβ in PBMC and ADH-THP-1 cells

PBMC undergoes spontaneous differentiation *in vitro* in liquid culture ^34^, as confirmed by Fig. 1 D,E. Thus, the spontaneous differentiation makes it difficult to explore the effect of 3D gel-like environment on PBMC. Therefore, we compared the response of chemical-inducers and 3D gel-like micro-environment on PBMC with that of differentiated THP-1 cells i.e ADH-THP-1. While we called the effect of inducers on monocytes (THP-1) as ‘primary response’, the effect of inducers on macrophages like PBMC/ADH-THP-1 is termed as ‘secondary response’. Fig. 8A depicts that 3D gel-like micro-environment causes a higher level of NF-κβ activation in ADH-THP-1 and PBMC cells. Like ADH-THP-1 cells, denial of adhesion by trypsin down regulates transcription of genes involved in adhesion (Fig. 8B). We also observe that providing a 3D gel cover to PBMC up-regulates the adhesion genes (Fig. 8B Col 1 vs Col 7 and Col 2 vs Col 8). Thus, the pathways shown in Fig. 6D is activated in PBMC by 3D gel-like micro-environment.

**Fig. 8.**
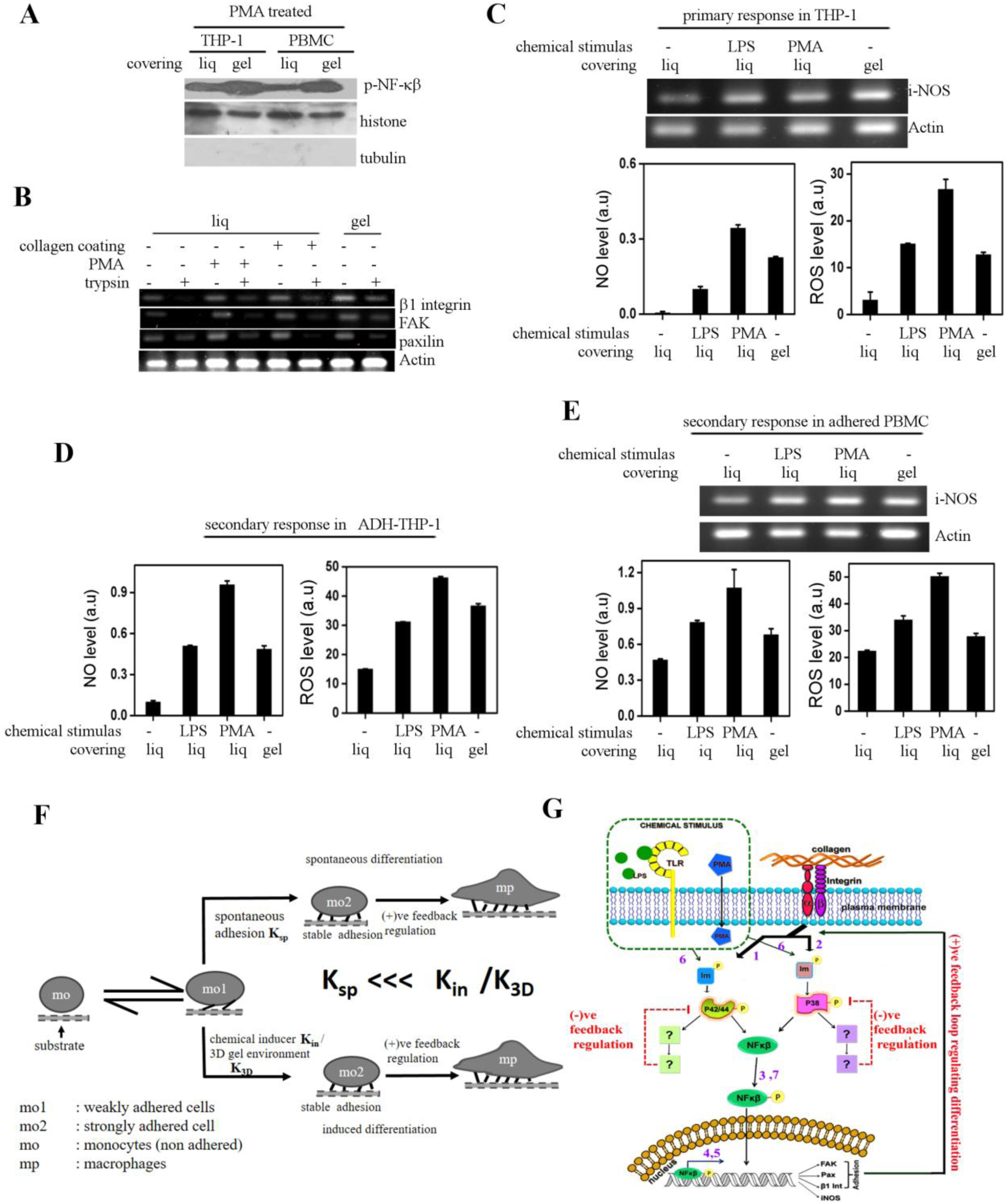
3-D gel-like micro-environment induces identical secondary response in PBMC and ADH-THP-1. (A) Western blot of p-NF-κβ in the nuclear fraction of PMA treated THP-1 and PBMC cells cultured with cover of identical chemical composition (0.1% agarose) in different physical state (liq: liquid, gel: agarose gel). (B) Semi-quantitative RT-PCR of adhesion genes β1-Integrin, Paxillin and FAK isolated from PBMC cells cultured under different condition in presence and absence of trypsin. (C) Semi-quantitative RT-PCR of i-NOS gene obtained 6 hrs post induction/seeding from THP-1 cells cultured on glass surface with 0.1% agarose liquid (liq)/gel cover on glass surface. The lower panels depict the quantitative comparison of NO (left) and ROS (right) production from NADH-THP-1 cells cultured in different conditions. (D-E) The quantitative comparison of NO (left) and ROS (right) production from 5 day old ADH-THP-1 (D) or adherent PBMC (E) 6 hrs post induction with LPS/PMA or 0.1% agarose liquid (liq)/gel covering on glass surface. Upper panel in ‘e’ depicts semi-quantitative RT-PCR of i-NOS obtained from 5 day old adherent PBMC in similar conditions as described previously. (F) Schematic representation depicting the relation between adhesion and differentiation for monocytes. (Mo: non-adhered monocytes. Mo1: weekly adhered monocytes, Mo2: strongly adhered monocytes, Mp: macrophages, K_sp_: Spontaneous rate of conversion of Mo1 to Mo2, K_in_ rate of conversion from Mo1 to Mo2 in response to chemical stimulation, K_3D_ rate of conversion from Mo1 to Mo2 in 3D gel-like micro-environment. (G) Cartoon representing the signaling cascade associated with the differentiation process mediated via chemical inducers and the adhesion. The Chemical inducers generate a short pulse of MAPK activation under negative feedback control. P-MAPK activates NF-κβ which translocates to the nucleus to transcribe adhesion genes. Stable adhesion complexes activates MAPK through a positive feedback loop to sustain higher levels of p-MAPK necessary for differentiation. The numbers in the cartoon represents the identification number of the arrow as used in table 1.

### 3D gel-like micro-environment elicit similar phenotypic response in PBMC and ADH-THP-1 cells

We also compared the phenotypic response of ADH-THP-1 and PBMC brought out by 3D gel cover. Our (Fig. 7I, Fig. 8C-E) and others study suggests an increase in NO and ROS production in monocytes (primary response) and differentiated macrophage (Fig. 8D secondary response) in response to LPS and PMA^35^ ^36,^. Interestingly, NADH-THP-1, ADH-THP-1, and adhered PBMC all of them show enhanced production of NO and ROS, in response to covering with agarose gel. The secondary response of PBMC is identical to that of ADH-THP-1 in 3D gel-like micro-environment (Fig. 8E). This suggests the significant influence of physical makeup of the micro-environment on PBMC, THP-1and HL60 cells.

**Table 1.** References for different links in the signalling network depicted in Fig. 8g

| Arrow. | Statement | Reference |
| --- | --- | --- |
| 1. | Integrin-mediated adhesion activates P42/44 MAPK | 37 |
| 2. | Integrin-mediated adhesion activates P38 MAPK | 38 |
| 3. | Integrin-mediated adhesion activates NF-κB | 30 |
| 4. | NF- κB involved in transcriptional upregulation of adhesion protein | 39 40 |
| 5. | NF- κB involved in transcriptional upregulation of differentiation-related genes | 41 33 42 |
| 6. | MAPK gets activated in monocyte in response to differentiation-inducing stimulus | 43 44 45 |
| 7. | NF- κB gets activated in monocyte in response to differentiation-inducing stimulus | 41 32 |

## Discussion

Various studies have explored the signaling associated with monocyte differentiation in response to chemical inducers in 2D cell culture system. However, these studies do not mimic the appropriate physicochemical micro-environment of the monocyte. Hence, fail to capture the effect of 3D gel-like micro-environments on differentiation. First, we establish that adhesion of monocytes is linked with its differentiation. Each monocyte has a small probability of undergoing spontaneous adhesion (K_sp_) as depicted in Fig. 8F (Fig. 1B,E). The adhesion initiates a positive feedback loop (Fig. 6D and Fig. 8G) to activate NF-κβ that triggers the differentiation. Treatment of chemical inducers in 2D fluid-like microenvironment activates a negative feedback loop on MAPK (Fig. 5A and Fig. 8G) to generate a short pulse of p-MAPK. Though. such a short pulse of p-MAPK is incapable of producing significant levels of p-NF-κβ, it is sufficient to induce adhesion by 1) transcriptional up-regulation of adhesion genes (Fig. 6A) and 2) by reducing the membrane fluctuation to facilitate adhesion in THP-1 and RAW cells (Fig. 4).As a result the probability of monocyte adhesion (K_in_/K_3D_)increases, which then activates the same positive feedback loop (Fig. 6D and Fig. 8G) to trigger the monocyte differentiation. The negative and positive feedback loop in Fig. 8G explains the necessity and sufficiency of adhesion in monocyte differentiation.

Our experimental method decouples the effect of a physicochemical micro-environment that arise out of its chemical makeup from that of its physical makeup. Such methods can be applied in other contexts to understand the influence of 3D gel-like micro-environment on other cellular functions. While the 3D gel-like micro-environment irrespective of its chemical makeup (agarose, alginate, matrigel, and collagen) is capable of triggering the positive feedback loop causing NF-κβ activation (Fig.:6D), identical chemical makeup in a liquid state is not capable of inducing differentiation via NF-κβ. We propose a signaling network for monocyte differentiation which explains the effect of chemical inducers, adhesion and 3D gel-like micro-environment each.

The human PBMC are pre-primed with cytokines/interleukins in the blood hence, most PBMC undergo differentiation. Yet, we do identify a small fraction of non-adherent PBMC just like NADH-THP-1 cells. Additionally, the 3D gel-like micro-environment increases the levels of p-NF-κβ in PBMC, ADH-THP-1 and THP-1cells. Thus, the behavior of PBMC is similar to that of the THP-1 cells. We also observe transcriptional up-regulation of i-NOS gene (Fig. 7I, Fig. 8C-E) involved in NO synthesis in 3D gel-like micro-environment. Since getting non-stimulated PBMC is difficult, we compared the secondary responses of PBMC in 3D gel-like micro-environment with that of PBMC in liquid-like micro-environment. Therefore, we establish that 3D gel-like micro-environment has a similar secondary response on PBMC and ADH-THP-1 cells as chemical stimulus. Thus, 3D gel-like micro-environment has a significant effect on the differentiation and activation of monocytes.

## Material and Methods

### Cell culture

Monocytic leukemia cell lines (THP-1 and HL-60) and RAW 264.7 cells of ATCC origin were obtained from Dr. Amitabho Sengupta and Dr.S.N. Bhattacharya from IICB. Both THP-1 and HL-60 were cultured in RPMI-1640 supplemented with 10% heat-inactivated FBS and 5% of Penstrep (Invitrogen and Himedia). RAW 264.7 cells were cultured in DMEM supplemented with 10% heat-inactivated FBS and 5% of Penstrep (Invitrogen and Himedia). THP-1 cells start to adhere within 4 to 6 hours post stimulation. At this time point cells are not dislodged by gentle aspiration but come out by washing. Human peripheral blood mononuclear cells (PBMC) were isolated by density gradient centrifugation protocol using ficol (Himedia) gradient. Blood samples from healthy human volunteers were collected in (anticoagulated 10% V/V) citrate buffer. The freshly collected blood samples layered over the top of ficol were centrifuged to obtain the buffy coat containing PBMC. The buffy coat was washed twice with PBS and subsequently dissolved in RPM1640, seeded into Petri dish that was incubated overnight. Next day, the suspended neutrophils and lymphocytes were washed with supplementation of fresh medium to the adhered PBMC.

Three distinct methodologies were employed to deny cellular adhesion to chemically stimulated THP-1 cells i) every 30 minutes after chemical stimulation the cells were mechanically dislodged from the plate (Fig. 3a), ii) stimulation on an adhesion incompatible substrate such as agar bed (Fig. 3b) and iii) culture in presence of trypsin on a glass bottom dish

### Microscopy and Image analysis

Cells were imaged with the sCMOS camera (Orca Flash 4.0, Hamamatsu) on an inverted fluorescence microscope from Carl Zeiss (Axio-observer Z1). Reflection Contrast Microscopy (RIM) was performed with an inverted fluorescence microscope (NIKON) equipped with EMCCD camera (Photometrics USA; evolve delta), using 60X 1.22 NA water immersion objective (additional 1.5X optical magnification was used). Image processing and analysis were done with ImageJ and Matlab. Total Internal Reflection Fluorescence Microscopy (TIRF) imaging was performed with an inverted fluorescence microscope (Olympus IX83) using 100X, 1.49NA with 488nm laser at a penetration depth of 70 nm, images were captured by sCMOS camera (Orca Flash 4.0, Hamamatsu).

### Phagocytosis assay

Carboxylated latex fluorescent beads of 200 nm (Invitrogen) were added to the cells. Two hours later the cells were washed with PBS to remove the floating beads. Fluorescence Z-stack images of the cells were acquired. Maximum intensity projection image from the previously acquired Z-stacks was obtained to count the total bead fluorescence per cell. The fluorescence intensity of z-projected images was scaled such that the background intensity becomes 1. The cells with average bead intensity more than 10% of the background were considered as phagocytotic.

### Single cell proliferation assay (SCPA)

The cells were treated with the chemical inducer for a given duration. Using serial dilution method, these cells were seeded in 96-well-plate such that most wells contained a single cell. Individual wells (having a single cell on day 0) were imaged to count the number of cells in them each day. The increase in cell number in any given well indicated proliferation hence no differentiation.

### MTT staining

Cells were washed with 1X PBS and incubated with MTT (Sigma) solution (1mg/ml) for 4 hours at 37 ^°^C. Subsequently the cells were lysed with DMSO and absorbance at 570 nm was measured.

### Genomic DNA isolation

Cells were lysed in lysis buffer (20 mM EDTA, 10 mM Tris pH 8.0, 200 mM NaCl, 0.2% Triton X-100, 100 mg/ml Pronase) by 1.5 hrs incubation at 37^°^C. The lysate was centrifuged at 14,000 rpm at room temperature (RT) for 5 minutes. The supernatant was mixed with equal volume of isopropanol and NaCl (such that its final concentration is 100 mM) for precipitation of DNA. The mixture was incubated overnight at - 20^°^C followed by centrifugation at 14,000 rpm at RT for 20-25 minutes to obtain the DNA pellet. The pellet was re-suspended in TE buffer and absorbance was measured at 260 nm.

### Cell culture with gel cover and isolation

THP-1 cells were suspended in the solution of 1% low melting agarose (Puregene, Hi res) in RPMI medium in a glass bottom Petri dish. The cells were allowed to settle down on the glass in agarose solution because of longer gelling time at 37^°^C. Fresh media was added after gelling of agarose. To isolate the cells, the medium was aspirated and agarose layer was peeled off the plate leaving behind the adhered cells.

### Cell culture in a 3D gel-like micro-environment

THP-1 cells were mixed with 0.1% low melting agarose solution in RPMI followed by 1-minute incubation at 4^°^C for quick jellification to prevent the cells from settling down on the glass. THP-1 cells were mixed with 1% sodium alginate (Sigma) solution in RPMI. The above suspension was added to 0.1M CaCl_2_ for jellification. After jellification, the CaCl_2_ solution was replaced with complete medium. For immobilization in a 3D collagen matrix, cells were mixed with collagen (BD Bioscience) in complete medium. 0.4% (V/V) 1N NaOH was added for collagen crosslinking. For immobilization in Matrigel, cells were mixed with diluted Matrigel (sigma) which get solidified at room temperature.

### Isolation of cells from the 3D micro-environment

Agarose matrix was molten by quick heating. Alginate/Collagen matrices were dissolved by treatment with 0.1M EDTA and 0.1N HCl respectively.

### Western blot

Cells were lysed with Laemmeli buffer. Prestained molecular markers (Biorad) were used to estimate the molecular weight of samples. Proteins were transferred to PVDF membranes (Millipore), after incubation with primary and secondary antibodies, bands were detected by ECL reagents.

### Immunostaining

Cells were fixed with 4% paraformaldehyde in PBS for 10 min, washed with PBS, blocked with 5% BSA for an hour. Primary antibody (Abcam) incubation was done overnight, washed with PBS and then incubated with Alexa tagged secondary antibody (CST) for 45 minutes, finally washed with PBS and imaged.

### FACS

Cells were fixed and permeabilized with 70% ethanol and cellular DNA was stained with propidium iodide (Sigma) and DNA amount was quantified in FACS.

### NO measurement

Griess reagent was incubated with spent medium and absorbance was measured at 540 nm.

### ROS measurement

25 µM DCFDA (Sigma) was incubated with spent medium for 30 minutes and the fluorescence was measured at Ex485 nm/Em535 nm.

### RT-PCR

RNA isolation was done using Trizol (Life-technologies). Then it was converted to cDNA. Subsequently desired gene was PCR amplified.

### Transfection of RAW cells

Cells were transfected using Fugene HD reagent (Promega) following manufactures protocol with Paxillin-GFP plasmid.

### Nucleus isolation

Equal number of cells from all conditions were lysed with ice-cold lysis buffer followed by centrifugation at 8000g for 15 minutes at 4°C to pellet down nuclei. Pellet was incubated with nuclear extraction buffer [20mM Tris-HCl (pH=7.9), 0.42M KCl, 0.2mM EDTA, 10% glycerol, 2mM DTT, 0.1mM PMSF and protease inhibitor cocktail] for 20 minutes at 4°C followed by centrifugation at 21,000 g for 15 minutes to precipitate nuclear debris. The lysates were processed further for analysis. Histone H3 was used as nuclear control protein.

## Acknowledgment

DKS acknowledges Grant No-SB/S0/BB-101/2013 (Department of Science and Technology, India), SR/S2/RJN-114/2011 (Ramanujan fellowship), BT/PR6995/BRB/10/1140/2012 (Department of Biotechnology, India). We thank Dr. Bidisha Sinha, IISER Kolkata for RIM and TIRF microscopy (funded by Grant no-IA/I/13/1/500885, Wellcome Trust-DBT India Alliance). We acknowledge CRNN, Kolkata for FACS and Dr. Dipyaman Ganguly, IICB Kolkata, for the PBMC culture facility and CSIR fellowship to AB (SPM) and MA.

## Additional information

PBMC were collected from informed healthy volunteer after taking proper consent as per the recommendation of IICB (Indian Institute for Chemical Biology, Kolkata) Research Ethics Board. All PBMC related work was done at IICB.

## Authorship Contribution

A.B and M.A. performed the experiments, analyzed the data and prepared the figures. R.M performed the experiments. P.S conceptualized the experiments. D.K.S. conceptualized the experiments analyzed the data and prepared the manuscript.

## Conflict of interest disclosure

The authors declare no competing financial interests.

## References

1. Wells, R. G. The role of matrix stiffness in regulating cell behavior. Hepatology 47, 1394–1400 (2008).

2. Cavo, M. et al. Microenvironment complexity and matrix stiffness regulate breast cancer cell activity in a 3D in vitro model. Sci. Rep. 6, (2016).

3. Gattazzo, F., Urciuolo, A. & Bonaldo, P. Extracellular matrix: a dynamic microenvironment for stem cell niche. Biochim. Biophys. Acta 1840, 2506–19 (2014).

4. Bloom, A. B. & Zaman, M. H. Influence of the microenvironment on cell fate determination and migration. Physiol. Genomics 46, 309–14 (2014).

5. Gordon, S. & Taylor, P. R. Monocyte and macrophage heterogeneity. Nat. Rev. Immunol. 5, 953–64 (2005).

6. Gordon, S., Plüddemann, A. & Martinez Estrada, F. Macrophage heterogeneity in tissues: Phenotypic diversity and functions. Immunol. Rev. 262, 36–55 (2014).

7. Shi, C. & Pamer, E. G. Monocyte recruitment during infection and inflammation. Nat. Rev. Immunol. 11, 762–774 (2011).

8. Tso, C., Rye, K.-A. & Barter, P. Phenotypic and Functional Changes in Blood Monocytes Following Adherence to Endothelium. PLoS One 7, e37091 (2012).

9. Ginhoux, F. & Jung, S. Monocytes and macrophages: developmental pathways and tissue homeostasis. Nat. Rev. Immunol. 14, 392–404 (2014).

10. Epelman, S., Lavine, K. J. & Randolph, G. J. Origin and functions of tissue macrophages. Immunity 41, 21–35 (2014).

11. Baieth, H. E. A. Physical parameters of blood as a non - newtonian fluid. Int. J. Biomed. Sci. 4, 323–329 (2008).

12. Nihat Özkaya, Margareta Nordin, David Goldsheyder, D. L. in Fundamentals of Biomechanics 86, 221–235 (Springer New York, 2012).

13. Swift, J. et al. Nuclear lamin-A scales with tissue stiffness and enhances matrix-directed differentiation. Science (80-.). 341, 1240104–15 (2013).

14. Handorf, A. M., Zhou, Y., Halanski, M. A. & Li, W. J. Tissue stiffness dictates development, homeostasis, and disease progression. Organogenesis 11, 1–15 (2015).

15. Basta, S., Knoetig, S., Summerfield, A. & McCullough, K. C. Lipopolysaccharide and phorbol 12-myristate 13-acetate both impair monocyte differentiation, relating cellular function to virus susceptibility. Immunology 103, 488–497 (2001).

16. Mittar, D., Paramban, R. & Mcintyre, C. Flow Cytometry and High-Content Imaging to Identify Markers of Monocyte-Macrophage Differentiation. BD Biosci. 1–19 (2011).

17. Hogg, N. et al. Identification of an anti-monocyte monoclonal antibody that is specific for membrane complement receptor type one (CR1). Eur. J. Immunol. 14, 236–243 (1984).

18. van Lochem, E. G. et al. Immunophenotypic differentiation patterns of normal hematopoiesis in human bone marrow: Reference patterns for age-related changes and disease-induced shifts. Cytometry 60B, 1–13 (2004).

19. Ayala, J. M. et al. Serum-induced monocyte differentiation and monocyte chemotaxis are regulated by the p38 MAP kinase signal transduction pathway. J. Leukoc. Biol. 67, 869–75 (2000).

20. Netea, M. G. et al. Interleukin-32 induces the differentiation of monocytes into macrophage-like cells. Proc. Natl. Acad. Sci. U. S. A. 105, 3515–20 (2008).

21. Schenk, M. et al. Interleukin-1β triggers the differentiation of macrophages with enhanced capacity to present mycobacterial antigen to T cells. Immunology 141, 174–180 (2014).

22. Traore, K. et al. Signal transduction of phorbol 12-myristate 13-acetate (PMA)-induced growth inhibition of human monocytic leukemia THP-1 cells is reactive oxygen dependent. Leuk. Res. 29, 863–79 (2005).

23. Aderem, A. Phagocytosis and the Inflammatory Response. J. Infect. Dis. 187, S340–S345 (2003).

24. Reddig, P. J. & Juliano, R. L. Clinging to life: cell to matrix adhesion and cell survival. Cancer Metastasis Rev. 24, 425–439 (2005).

25. Murphy, J. A., Franklin, T. B., Rafuse, V. F. & Clarke, D. B. The neural cell adhesion molecule is necessary for normal adult retinal ganglion cell number and survival. Mol. Cell. Neurosci. 36, 280–292 (2007).

26. Somaiah, C. et al. Collagen promotes higher adhesion, survival and proliferation of mesenchymal stem cells. PLoS One 10, 1–15 (2015).

27. Rao, K. M. K. MAP kinase activation in macrophages. jleukbio 69, 3–10 (2015).

28. Limozin, L. & Sengupta, K. Quantitative reflection interference contrast microscopy (RICM) in soft matter and cell adhesion. ChemPhysChem 10, 2752–2768 (2009).

29. Klein, K., Rommel, C. E., Hirschfeld-Warneken, V. C. & Spatz, J. P. Cell membrane topology analysis by RICM enables marker-free adhesion strength quantification. Biointerphases 8, 1–13 (2013).

30. Chen, J. et al. αvβ3 Integrins Mediate Flow-Induced NF-κB Activation, Proinflammatory Gene Expression, and Early Atherogenic Inflammation. Am. J. Pathol. 185, 2575–2589 (2015).

31. Center, R. R., Roche, H. & Collec-, C. Regulation of Adhesion and Growth of Fibrosarcoma Cells by NF-KB RelA Involves Transforming Growth Factor 13. Mol Cell Biol. 14, 5326–5332 (1994).

32. Holden, N. S. et al. Phorbol ester-stimulated NF-κB-dependent transcription: Roles for isoforms of novel protein kinase C. Cell. Signal. 20, 1338–1348 (2008).

33. Lowe, J. M. et al. p53 and NF-κB coregulate proinflammatory gene responses in human macrophages. Cancer Res. 74, 2182–92 (2014).

34. Zhang, M. & Huang, B. The multi-differentiation potential of peripheral blood mononuclear cells. Stem cell Res. & Ther. 3, 48 (2012).

35. Kuwabara, W. M. T. et al. NADPH oxidase-dependent production of reactive oxygen species induces endoplasmatic reticulum stress in neutrophil-like HL60 cells. PLoS One 10, 1–15 (2015).

36. Sekhar, S., Sampath-Kumara, K. K., Niranjana, S. R. & Prakash, H. S. Attenuation of reactive oxygen/nitrogen species with suppression of inducible nitric oxide synthase expression in RAW 264.7 macrophages by bark extract of Buchanania lanzan. Pharmacogn. Mag. 11, 283–291 (2015).

37. Yee, K. L. & Hammer, D. A. Integrin-mediated signalling through the MAP-kinase pathway. IET Syst. Biol 2, 8–15 (2008).

38. Aikawa, R., Nagai, T., Kudo, S., Akazawa, H. & Issei, K. Integrins play a critical role in mechanical stress-induced p38 activation. Hypertension 39, 233–238 (2002).

39. Zhang, L. L. et al. Phosphatase and tensin homolog (PTEN) represses colon cancer progression through inhibiting paxillin transcription via PI3K/AKT/NF-κB pathway. J. Biol. Chem. 290, 15018–15029 (2015).

40. Ahmed, K. M., Zhang, H. & Park, C. C. NF-B Regulates Radioresistance Mediated By 1-Integrin in Three-Dimensional Culture of Breast Cancer Cells. Cancer Res. 73, 3737–3748 (2013).

41. Chen, B. C. & Lin, W. W. PKC- and ERK-dependent activation of I kappa B kinase by lipopolysaccharide in macrophages: enhancement by P2Y receptor-mediated CaMK activation. Br. J. Pharmacol. 134, 1055–1065 (2001).

42. Morgan, M. J. & Liu, Z. Crosstalk of reactive oxygen species and NF-κB signaling. Cell Res. 21, 103–15 (2011).

43. Guha, M. et al. Lipopolysaccharide activation of the MEK-ERK1/2 pathway in human monocytic cells mediates tissue factor and tumor necrosis factor alpha expression by inducing Elk-1 phosphorylation and Egr-1 expression. Blood 98, 1429–39 (2001).

44. Saldeen, J. & Welsh, N. p38 MAPK inhibits JNK2 and mediates cytokine-activated iNOS induction and apoptosis independently of NF-αα translocation in insulin-producing cells. Eur. Cytokine Netw. 15, 47–52 (2004).

45. van der Bruggen, T., Nijenhuis, S., van Raaij, E., Verhoef, J. & van Asbeck, B. S. Lipopolysaccharide-induced tumor necrosis factor alpha production by human monocytes involves the raf-1/MEK1-MEK2/ERK1-ERK2 pathway. Infect. Immun. 67, 3824–9 (1999).

